# Mechanism of PMC (2,2,5,7,8-pentamethyl-6-chromanol), a sterically hindered phenol antioxidant, in rescuing oxidized low-density lipoprotein-induced cytotoxicity in human retinal pigment epithelial cells

**DOI:** 10.1101/2025.06.19.660627

**Authors:** Suman Chaudhary, Jean Moon, Zhengping Hu, Emil Kriukov, Sergio Pestun, Petr Y Baranov, Yin-Shan Eric Ng, Patricia A. D’Amore

## Abstract

Geographic atrophy or late stage dry age-related macular degeneration (AMD) is characterized by drusen deposition and progressive retinal pigment epithelium (RPE) degeneration, leading to irreversible vision loss. The formation of drusen leads to dyshomeostasis, oxidative stress and irreversible damage to RPE. In this study, we used an in vitro model of oxidized-low density lipoproteins (ox-LDL) induced human RPE damage/death model to investigate the mechanism whereby a sterically hindered phenol antioxidant compound, PMC (2,2,5,7,8-pentamethyl-6-chromanol) protects RPE against ox-LDL-induced damage. We show that PMC exerts its protective effect by preventing the upregulation of stress-responsive heme oxygenase-1 (HMOX1/HO-1) and NAD(P)H:quinone oxidoreductase (NQO1) at mRNA and protein levels. This effect was due to PMC’s blockade of ROS generation, which in turn blocked nuclear translocation of the Nuclear factor erythroid 2-related factor 2 (Nrf2) transcription factor, ultimately preventing the upregulation of antioxidant response elements (ARE), including HMOX1 and NQO1. A key role for HO-1 was demonstrated when the protective effect of PMC was inhibited by the knockdown of HMOX1. Additionally, treatment of PMC under different experimental conditions and time points revealed that the continuous presence of PMC is required for optimal protection against ox-LDL-induced cytotoxicity, defining the cellular pharmacokinetics of the molecule. Our data demonstrate the involvement of a key antioxidant pathway through which PMC mitigates oxidative stress induced by ox-LDL and provides a potential therapeutic strategy to suppress RPE degeneration/damage during AMD progression.

## Introduction

Late-stage dry age-related macular degeneration (AMD) also known as geographic atrophy (GA) is a degenerative disease of the retina. It is characterized by slow and irreversible damage of the RPE, photoreceptors, and choriocapillaris, leading to central visual impairment^1^. In the retina, triglycerides and cholesterol are important sources of lipid metabolism. However, due to aging and genetic variants, there is dysregulated lipid metabolism that can contribute to the deposition of lipoproteins and lipid rich drusen between the RPE and Bruch’s membrane ^2^. Additionally, owing to the high oxygen metabolic environment in the retina and the resulting significant generation of reactive oxygen species (ROS), the retina is highly susceptible to oxidative stress. This oxidative stress has often been linked with lipid peroxidation, endoplasmic reticulum stress, chronic inflammation and DNA damage, which all contribute to the development of macular degeneration^3-5^. In particular, oxidative stress and accumulation of oxidized lipids and lipoproteins exacerbated by dysregulated lipid metabolism have been linked with RPE dysfunction, degeneration, and death^6-8^.

Typical antioxidant enzymes reduce the ROS levels by scavenging free radicals, restoring the cellular redox status to promote repair of cellular damage. However, excessive pro-oxidative conditions such as aging, cigarette smoke, high-fat diet, and lack of antioxidant rich diet can alter redox homeostasis, leading to retinal degeneration^9^. Previously, a multi-center randomized controlled Age-Related Eye Disease Studies (AREDS) and AREDS 2 have indicated the efficacy of antioxidant supplementation in reducing the progression of dry AMD^10,11^ which was recently confirmed by a follow-up post hoc analysis study^11,12^, thus, highlighting the importance of antioxidant compounds in reducing AMD pathology.

One of the primary pathways employed by RPE cells to mitigate oxidative stress and maintain cellular homeostasis is the nuclear factor E2-related factor 2 (Nrf2) antioxidant response element (ARE) pathway. Upon oxidative stress, dissociation of Nrf2 from its cytoplasmic anchor Kelch-like ECH-associated protein 1 (Keap1) mediates its nuclear translocation and the subsequent transcriptional upregulation of targets of the ARE pathway ^13-15^. Among the component of this pathway, two key antioxidant enzymes; heme oxygenase-1 (HO-1) and NAD(P)H: quinone oxidoreductase (NQO1) have been shown to protect human RPE cells against oxidative stress by counteracting ROS^16,17^. HO-1, also known as heat shock protein 32 (HSP32), is involved in the degradation of heme which leads to the production of carbon monoxide (CO), ferrous iron, and biliverdin products. The expression of HO-1, encoded by the HMOX1 gene, is regulated at both transcriptional and translational levels^18^. It is a pro-oxidant indicator, as it is induced by a broad range of factors such as cytokines, hormones and growth factors, tissue damage, and exposure to blue light^19-21^ in the retina. HO-1 is involved in various pathological processing including oxidative stress, inflammatory response, and ferroptosis^20^. It is an important antioxidant and a crucial component of cellular defense system. However, studies have documented its dual nature where overexpression of HO-1 is linked to excessive ROS generation. Numerous recent studies have indicated the upregulation of HO-1 in disease models of AMD, including in vitro systems using blue light^22^ and in vivo models employing sodium iodate^23^ and light exposure ^24^. Studies have also shown that high levels of a free ferrous ion (Fe^2+^), a catabolic product of surplus HO-1, can cause lipid peroxidation and oxidative stress exacerbated by ferroptosis. This occurs mainly due to the vicious cycle between the upregulation of HO-1 and oxidative stress resulting in iron overload^20,23^. Therefore, it is imperative to maintain ambient Nrf2/HO-1 balance for effective therapeutics.

We have previously developed an oxidized-low density lipoproteins (ox-LDL)-induced human RPE injury in vitro model that involves the induction of ROS generation and lysosomal destabilization, and ultimately RPE damage/death and AMD-associated pathogenesis^25^. Utilizing this model, we have identified a class of sterically hindered phenols with free radical scavenging properties that robustly protects against ox-LDL induced RPE damage and cell death. PMC (2,2,5,7,8-pentamethyl-6-chromanol), a prototypic member of this family, also has the ability to reduce the levels of intracellular cholesterol through the NPC-1/NPC-2 efflux pathway^26^. However, the underlying molecular mechanism of the protective PMC activity against ox-LDL induced RPE damage is not yet fully understood and may include scavenging of ROS and other direct and indirect antioxidant activities. The goal of this current study is to elucidate the antioxidant mechanism involved in PMC-mediated protection of RPE cells from ox-LDL–induced cytotoxicity.

## Methods

### Cell culture

Primary human fetal retinal pigmented epithelium cells (hRPE) (Lonza, USA) were cultured in 50% (v/v) DMEM/F12 (Life Technologies, USA)/50% (v/v) αMEM (Sigma-Aldrich, USA) supplemented with 1x penicillin-streptomycin, Glutamax, sodium pyruvate, non-essential amino acids (all from Life Technologies, USA). Additionally, 10% (for growth media) or 2% (for maturation media) heat-inactivated fetal bovine serum (FBS) (Sigma-Aldrich, USA), 10 mM, 50% N1 media supplement, 0.25 mg/ml taurine, 0.02 μg/ml hydrocortisone, and 0.013 ng/ml 3,3,5-triiodo-L-thyronine (all from Sigma-Aldrich, USA) were added in accordance to the previously published protocol^27^. Half of the media was replenished every 2-3 days for 4 weeks to ensure maturation and pigment accumulation in hRPE cells. Cells were then incubated in serum-free media supplemented overnight prior to treatments. Human ARPE-19 cells (American Type Culture Collection, Manassas, USA) were cultured in DMEM/F-12 media with L-glutamine supplemented with 1x penicillin-streptomycin and 10% FBS.

### Oxidized-LDL (ox-LDL) and PMC treatments

Primary hRPE or ARPE-19 cells were cultured at a seeding density of 0.3×10^6^/well for 6-well plates for 2 to 4 weeks, incubated in serum-free overnight and then treated with 200 µg/ml oxidized-LDL (ox-LDL) (cat no-770252-4, Kalen Biomedical, LLC, USA) with or without 1.3 µM PMC (cat no-430676, Sigma-Aldrich, USA) for 24 and 48 hr. DMSO alone (PMC diluent) was used as a control. Conditioned media were collected for analysis of cytotoxicity, while RNA isolation or lysate from the treatment groups were collected for subsequent studies.

### Cytotoxicity assay

LDH assay (cat no-4744934001, Sigma-Aldrich, USA) was conducted to quantify cytotoxicity. Conditioned media from cells treated with 1% Triton X-100 were used as positive controls with maximum LDH activity. Spontaneous LDH release was measured from control cells without ox-LDL and PMC treatments. Percentage cytotoxicity was estimated using the formula.

% cytotoxicity= [Experimental LDH activity-spontaneous LDH activity]/ maximum LDH activity-spontaneous LDH activity]×100.

### Bulk RNA sequencing

Total RNA was extracted using RNeasy plus mini kit (Cat no-74034, Qiagen, USA) from all the treatment groups of hRPE cells, in accordance with the manufacturer’s instructions. The concentration and quality were determined using the NanoDrop Spectrophotometer. Bulk RNA sequencing was conducted by Azenta Life Sciences, USA and sequencing platform used was Illumina^@^NovaSeq^TM^. Sequence reads were trimmed using Trimmomatic V.0.26 to remove nucleotides with poor quality and possible adaptor sequences. The trimmed reads were mapped to Homo sapiens GRCh38 reference genome using the STAR aligner v.2.5.2b. Unique gene hit counts were estimated using feature counts from the subread package v.1.5.2, following which differential gene expression was analyzed. Wald test was used to generate p-values and log2fold changes. Genes with adjusted p-value of <0.05 and log2fold change of >1 were termed as differentially expressed genes for each comparison. Gene ontology analysis was conducted on the statistically significant genes using the software GeneSCF V.1.1-p2. The DESeq2 (https://genomebiology.biomedcentral.com/articles/10.1186/s13059-014-0550-8) output was used for the conditions of interest, and the GSEA (Gene Set Enrichment Analysis) approach was applied for every treatment group vs. Control independently.

Data availability: The code generated in this study is deposited on GitHub and available by the following link: https://github.com/mcrewcow/Suman_bulkRNAseq/tree/main. DESeq2, GSEA, and the pathways-to-patterns transformation data are available on GitHub.

### Quantitative PCR

RNeasy plus mini kit (Cat no-74034, Qiagen, USA) was used to extract total RNA from the different treatment groups of the hRPE and ARPE-19 cells and the quality (A260/A280: ∼2.0) and concentration (1µg) of RNA was estimated using NanoDrop Spectrophotometer (Thermo Fisher Scientific, USA). cDNA synthesis was conducted using the SuperScript IV VILO Mastermix (cat no-11756050, Life Technologies, USA) as per the manufacturer’s instructions. Techne TC-512 Thermal Cycler (Thermo Fisher Scientific, USA) was used for annealing primer (25°C, 10 min), reverse transcription (50°C for 10 min) and inactivation of enzyme (85°C for 5 mins). Real time qPCR was conducted using the PowerUp SYBR Green Master Mix (cat no-A25776, Thermo Fisher Scientific, USA) using LightCycler 480 (Roche, Switzerland). Thermal cycling conditions of 50°C for 2 min, 95°C for 2 mins, 40 cycles of 95°C for 15 sec and 60°C for 1 min was performed in duplicate, three times for each treatment group. Amplification specificity was confirmed using the melt curve analysis. Ct values were normalized to Gapdh (housekeeping gene), and analysis was performed in comparison to the control group. List of primer sequences are provided in **Table 1**.

**Table 1.**
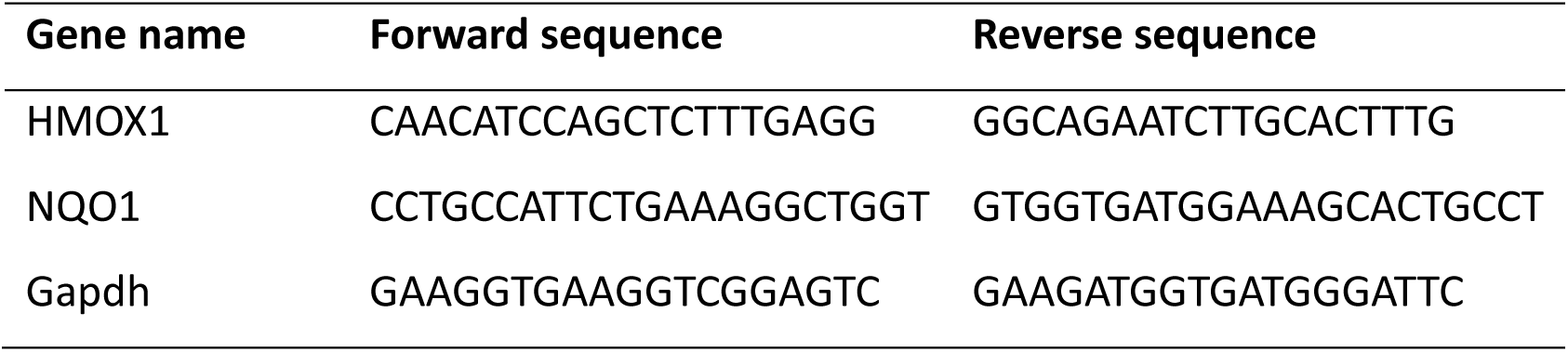
Primer sequences.

### siRNA gene knockdown

HRPE cells were seeded at a density of 0.3×10^6^/well on six-well plates and maintained for 2 to 4 weeks. For transfection, siRNA for HMOX1 (cat no-sc-35554, Santa Cruz Biotechnology, USA) consisting of 3 target-specific 19-24nt siRNAs and scrambled siRNA control (cat no sc-37007, Santa Cruz Biotechnology, USA) were added into hRPE cells using lipofectamine RNAiMAX (cat no-13778075, Thermo Fisher Scientific, USA) as per the manufacturer’s instructions. Following transfection for 48 hr, the scrambled siRNA and siRNA HMOX1 transfected cells were treated with ox-LDL and ox-LDL along with PMC for 24 or 48 hr. Conditioned media were collected from each treatment group to conduct cytotoxicity assay for each treatment group and time point using LDH kit. Lysates were collected for immunoblotting analysis.

### Western blot

Primary hRPE or ARPE-19 cells were lysed by adding 1X cell lysis buffer (cat no-9803s, Cell Signaling Technology, USA) containing protease inhibitor (cat no-5871S, Cell Signaling Technology, USA) to the cell pellet for 20 min with occasional vortexing, followed by centrifugation at 12,000 × g for 20 minutes at 4°C. The supernatant containing total protein was collected, and protein concentration was determined using the BCA assay. Equal amount of protein (30µg) was loaded onto a 4-20% SDS-PAGE gel and fractionated by electrophoresis. Following this, the proteins were transferred onto a nitrocellulose membrane. The membranes were blocked in 3% bovine serum albumin for 1 hr at room temperature and incubated overnight with primary antibodies listed in **Table 2**. After washing with Tris buffered saline with Tween 20 (TBST), the membrane was incubated with secondary antibody and further washed with TBST. The fluorescent band intensity in membranes were imaged using infrared imaging system (Odyssey CLx Imaging System; LI-COR Biosciences). Band intensities were quantified using ImageJ 1.53ts software (NIH) and graphically analyzed using GraphPad Prism (GraphPad Software Inc, USA), version 10.0.3 after normalizing with the housekeeping protein (GAPDH or β-tubulin).

**Table 2.** List of antibodies.

| Antibody | Host species* | Species reactivity* | Company | Catalogue number | Dilution** |
| --- | --- | --- | --- | --- | --- |
| HO-1 | m | b, h, m, r, d | Thermo<br>Fisher<br>Scientific | MA1-112 | WB: 1:1000<br>ICC: 1:200 |
| NQO1 | m | h | Abcam | ab28947 | WB: 1:500 |
| Calreticulin | Rb | h, m, r | Cell Signaling<br>Technology | D3E6 | ICC: 1:200 |
| Nrf2 | m | h | Santa Cruz<br>Biotechnology | sc-365949 | WB: 1:500<br>ICC: 1:100 |
| Gapdh | Rb | h, m, r, mk | Cell Signaling<br>technology | D16H11 | WB: 1:1000 |
| α-tubulin | Rb | h, m, r, mk,<br>b | Cell Signaling<br>Technology | 2144 | WB: 1:1000 |
| * m- mouse, Rb- rabbit, h- human, r- rat, b-bovine; d, dog, mk-monkey |  |  |  |  |  |
| **WB-western blotting, ICC- immunocytochemistry |  |  |  |  |  |

### Immunocytochemistry

ARPE-19 cells were seeded at a density of 7.5 x 10^4^ cells on 12 mm coverslips for 2 weeks as previously described^26^. The cells were then incubated overnight in serum-free media and treated with the ox-LDL with or without PMC for 24 to 48 hr. Cells were fixed using 4% paraformaldehyde for 20 min and permeabilized using 0.1% Triton X-100 then blocked for 1 hr with 3% bovine serum albumin and incubated overnight with mouse anti-HO-1, rabbit anti-calreticulin or mouse anti-Nrf2 antibody (**Table 2**). Next, the cells were incubated with Alexa Fluor 488-labeled donkey anti-rabbit (1:500, A-21206, Thermo Fisher Scientific, USA), Alexa Fluor 594-labeled donkey anti-mouse (1:500, A-21203, Thermo Fisher Scientific, USA) or Alexa Fluor 488-labeled donkey anti-mouse (1:500, A-21202, Thermo Fisher Scientific, USA). The cells were mounted with ProLong gold antifade mounting media with DAPI (cat no- P36935, Life Technologies, USA) and imaged using a Axioscope 2 Mot plus microscope (Carl Zeiss Meditec AG, Zeiss, Germany). Nrf2 nuclear translocation was imaged using SP8 confocal microscope (Leica, Germany). Multiple focal planes were imaged using Z-stacking in the confocal microscope to create a three-dimensional image. For each experiment, a representative image from a minimum of six different fields was acquired. Colocalization was quantified using Manders’ coefficient in the JaCoP plugin of ImageJ 1.53ts software (NIH) to measure the proportion of overlap between the two fluorescence where 0 indicates no overlap while 1 indicates complete overlap.

### Reactive oxygen species (ROS) detection

For ROS detection, ARPE-19 cells were cultured on 12 mm coverslips at a density of 7.5 x 10^4^ cells for 2 weeks. Cells were incubated in serum-free media overnight and treated with control (DMSO diluent), ox-LDL (100µg) with or without PMC (1.3 µM) and PMC alone for 24 hr. Cells were then incubated with 1 µM of H2DCFDA (H2-DCF, DCF) (cat no-D399, Invitrogen, USA) for 30 min. For endoplasmic reticulum specific staining, cells were incubated with 1 µM of ER-Tracker™ Red (BODIPY™ TR Glibenclamide) (cat no- E34250, Invitrogen, USA) for 20 min at 37°C then fixed with 4% paraformaldehyde for 20 min. Hoechst 33342 was used to stain the nuclei. Visualization of cells was accomplished using the Axioscope 2 Mot plus microscope (Carl Zeiss Meditec AG, Zeiss, Germany). Colocalization quantification was performed using Manders’ coefficient in the JaCoP plugin of ImageJ 1.53ts software (NIH).

### Statistical analysis

The results are presented as mean ± SEM of a minimum of three biological replicates from three independent experiments. Statistical significance was measured using one-way ANOVA followed by Tukey’s multiple comparison test, using GraphPad Prism version 10.0.3 between the control and treatment groups. Values of p < 0.05 were considered as statistically significant.

## Results

### PMC protects RPE against ox-LDL induced cytotoxicity

Challenge with ox-LDL on primary hRPE cells resulted in significant cell death of 27.9 ± 0.8% at 24 hr and 44.9 ± 1.3% at 48 hr compared to control using LDH assay. Inclusion of PMC led to a significant (p< 0.001) reduction in cell toxicity to 17.2 ± 0.6% and 8.6 ± 1.3% at 24 and 48 hr, respectively, in comparison to control. PMC (1.3 µM) alone did not affect cell viability compared to control at 48 hr (Figure 1A).

**Figure 1.**
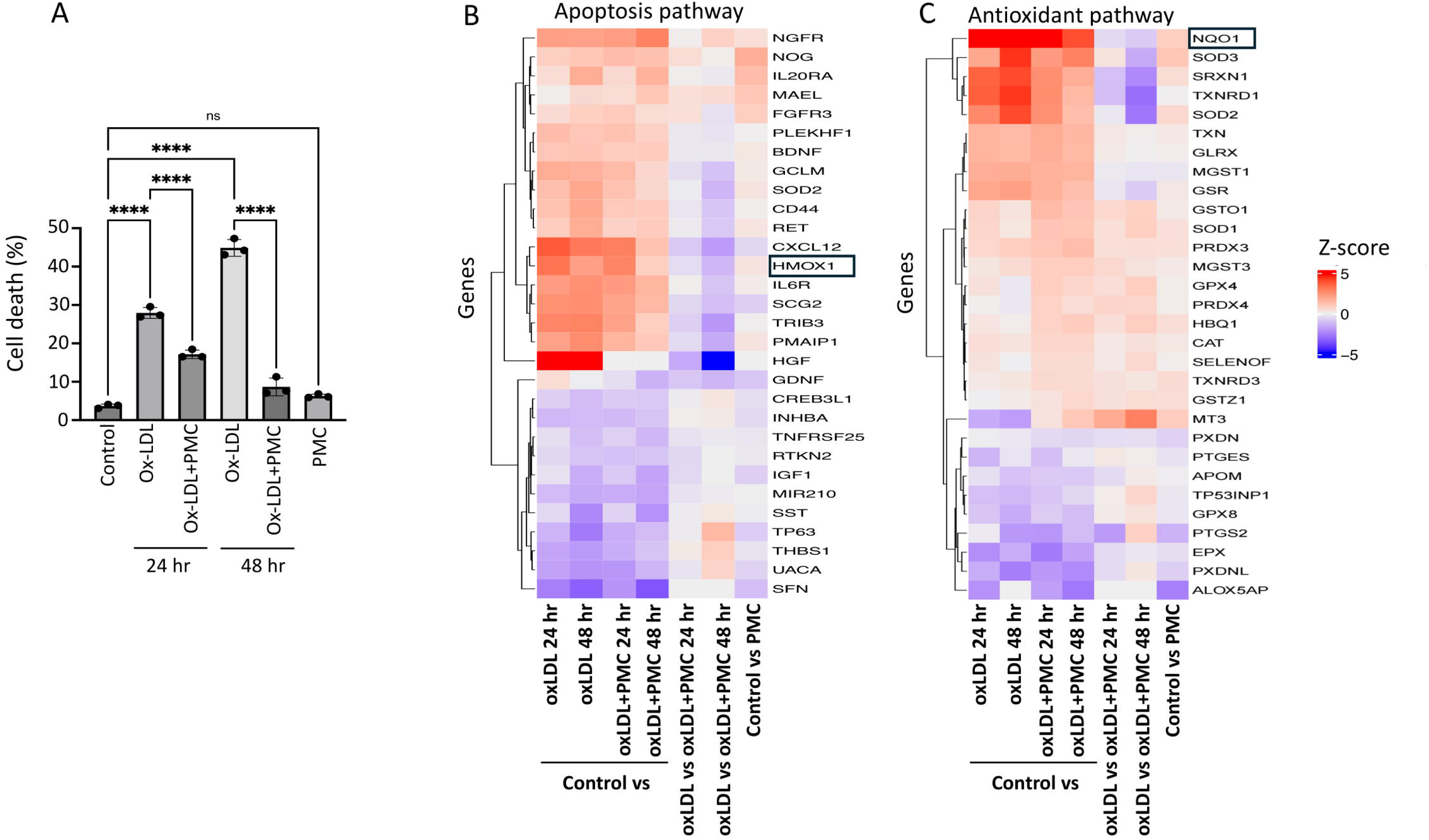
PMC protects hRPE cells from ox-LDL-induced damage by modulating apoptotic and antioxidant pathways. (A) hRPE cells were treated with 200 µg/ml for 24 and 48 hr in the absence or presence of PMC (1.3 µM). Cytotoxicity was also measured in untreated (control) and PMC alone treated cells. Cell death was measured in the condition media using the LDH assay, n=3. Values are expressed as mean ± SEM. Statistical analysis was conducted using one-way ANOVA, ***P<0.001, n=3. Heatmap of top differently expressed genes of bulk RNA sequencing in (B) apoptosis and (c) antioxidant pathways, n=3. Colors shows intensity in z-scored units where red shows replicates with high expression (z-score= +5) and blue shows replicates with low expression (z-score =-5).

To gain insights into the mechanisms involved in PMC-mediated protection of hRPE cells from ox-LDL-induced toxicity, bulk RNA sequencing was conducted. A heatmap of the differentially expressed genes between control vs. ox-LDL, ox-LDL and PMC (24 and 48 hr), PMC alone, and ox-LDL vs. ox-LDL and PMC (24 and 48 hr) indicated modulation of genes involved in the apoptosis and antioxidant pathways (Figure 1B, C). Interestingly, there was an upregulation of antioxidant response element (ARE) genes HMOX1 and NQO1 in cells treated with ox-LDL at 24 and 48 hr in comparison to control. On the other hand, treatment with PMC and ox-LDL led to significantly lower levels of HMOX1 and NQO1 compared to cells treated with ox-LDL alone at 24 and 48 hr (Figure 1B, C).

### ox-LDL upregulates heme oxygenase-1 in RPE cells

Having identified the differentially expressed genes that are potentially involved in PMC mediated protection of hRPE cells against ox-LDL, we validated the RNA sequencing results at transcriptional and translational levels. Semi-quantitative qPCR analysis revealed a significant increase in HMOX1 (encodes for the HO-1 protein) mRNA in comparison to control at both 24 and 48 hr upon ox-LDL challenge. Treatment with ox-LDL and PMC led to a significant reduction of ox-LDL-induced HMOX1 mRNA level. At 24 hr, an ox-LDL challenge led to a 307.5 ± 25.2-fold increase in HMOX1 compared to control, whereas treatment with PMC together with ox-LDL resulted in a significantly lower 26.2 ± 11.0-fold increase in HMOX1 compared to control (p <0.05, ox-LDL vs. ox-LDL+PMC). At 48 hr, ox-LDL alone induced a 382.3 ± 135.0-fold increase in HMOX1 expression compared to control while ox-LDL challenge with PMC treatment resulted in a significant lower increase in HMOX1 expression at 74.3 ± 31.7-fold compared to control (p <0.05 ox-LDL vs. ox-LDL+PMC). PMC alone had no effect on HMOX1 levels in comparison to control (Figure 2A).

**Figure 2.**
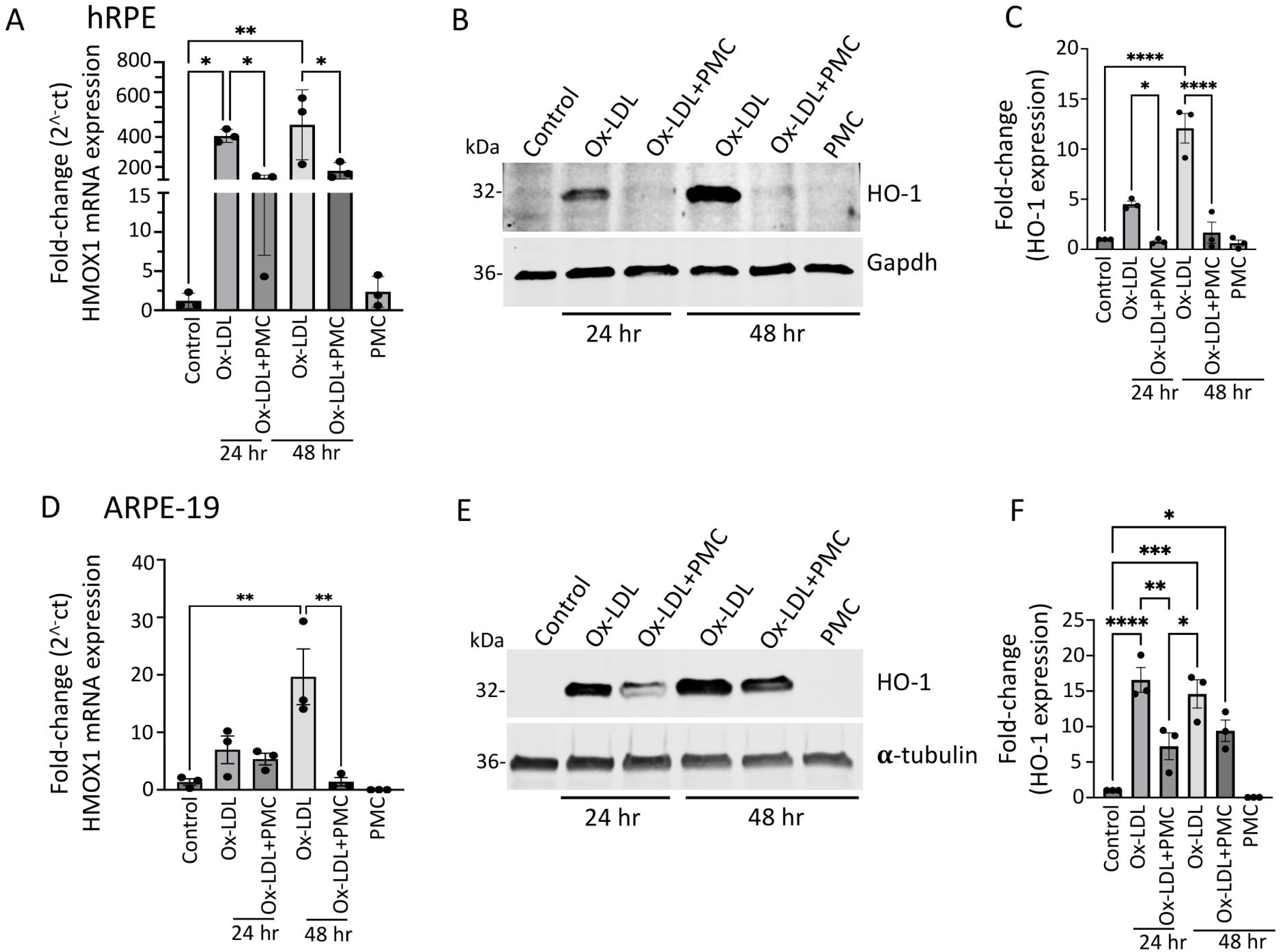
Heme oxygenase-1 upregulation in ox-LDL-treated RPE cells is suppressed by treatment with PMC. (A) hRPE cells matured for 4 weeks, were treated with ox-LDL (200 µg/ml) with or without the presence of PMC (1.3 µM) in serum-free media. Relative HMOX1 transcript levels were measured using qPCR. (B) Western blotting of the hRPE cells with the above indicated treatments at 24 and 48 hr was conducted to examine HO-1 levels. (C) Quantification was performed using densitometry after normalization with GAPDH, n=3. (D) HMOX-1 mRNA levels were determined by PCR in serum-starved ARPE-19 cells treated with ox-LDL in the presence or absence of PMC and qPCR. (E) Cell lysates from ARPE-19 cells treated as above-were examined using western blot to determined HO-1 levels. (F) Quantification of HO-1 levels were conducted after normalization with **α**-tubulin. HMOX1/HO-1 was also assessed in hRPE and ARPE-19 cells treated with PMC alone. Serum-starved untreated cells were considered as control for all the experiments. Values are indicated as mean ± SEM of n=3. One-way ANOVA was used for statistical analysis, *P<0.05, **P<0.01, ***P<0.001

Western blotting analysis of hRPE revealed a low expression of HO-1 protein in control and PMC treated cells (Figure 2B and C). However, there was a significant increase in the levels of HO-1 protein in cells treated with ox-LDL at 24 and 48 hr. Quantification showed a 4.5 ± 0.3-fold increase in HO-1 levels at 24 hr in comparison to untreated control, which was significantly reduced to 0.79 ± 0.1-fold in cells treated with ox-LDL and PMC (p<0.05, ox-LDL vs. ox-LDL+PMC). At 48 hr, HO-1 levels were significantly increased by 12.1 ± 1.5-fold (p<0.001) in cells treated with ox-LDL in comparison to control, while addition of PMC drastically suppressed the ox-LDL-induced upregulation of HO-1 protein to 1.7 ± 1.1 (p<0.001 ox-LDL vs. ox-LDL+PMC) (Figure 2C).

Similar results were observed in ARPE-19 where there was a 7.0 ± 2.4-fold increase in HMOX1 mRNA levels at 24 hr and 19.6 ± 4.9-fold increase at 48 hr in cells treated with ox-LDL comparison to control. The addition of PMC reduced HMOX1 levels to 5.4 ± 1.0-fold increase at 24 hr. At 48 hr addition of PMC resulted in a 1.4 ± 0.7-fold increase in HMOX1 levels compared to untreated control, which is significantly lower compared to ox-LDL alone (p<0.01 ox-LDL vs. ox-LDL+PMC). PMC alone had no effect on HMOX1 mRNA expression (Figure 2D). Western blot analysis corroborated these results. Control cells showed low HO-1 protein expression, and PMC had no effect on HO-1 protein levels. The ox-LDL challenge resulted in upregulation of HO-1 at 24 and 48 hr, which were reduced by the addition of PMC (Figure 2E). Quantification showed that ox-LDL induced a significant increase in HO-1 protein at 24 hr (16.6 ± 1.8-fold; p<0.001) and 48 hr (14.6 ± 1.9-fold; p<0.001) compared to control. PMC addition with ox-LDL at 24 hr resulted in a significantly lower the HO-1 level (7.2 ± 1.9-fold) in comparison to ox-LDL alone (p<0.01, ox-LDL vs ox-LDL+PMC), and at 48 hr HO-1 protein levels were also trending lower (9.4 ± 1.5-fold compared to control) compared to ox-LDL alone, but the data were not statistically significant (Figure 2F).

### ox-LDL upregulates NQO1 in RPE, which is lowered by PMC

ox-LDL treatment led to a significant upregulation of NQO1 mRNA levels at 24 hr (6.5 ± 0.7-fold; p<0.001) and 48 hr (4.0 ± 0.4-fold) in comparison to controls, which was significantly reduced by PMC at 24 hr (3.7 ± 0.4-fold; p<0.001, ox-LDL vs. ox-LDL+PMC) and trending a reduction at 48 hr (2.4 ± 0.5-fold) that was not statistically significant (Figure 3A). Western blot analysis indicated a similar trend in NQO1 protein expression; a 2.8 ± 0.3-fold increase in NQO1 levels was observed in ox-LDL treated cells at 24 hr in comparison to control levels, which were decreased to 1.6 ± 0.2-fold by the addition of PMC. At 48 hr, both ox-LDL alone and ox-LDL+PMC treated cells showed similar upregulation of NQO1 levels at 3.0 ± 0.5-fold and 2.8 ± 0.4-fold, respectively (Figure 3B, C).

**Figure 3.**
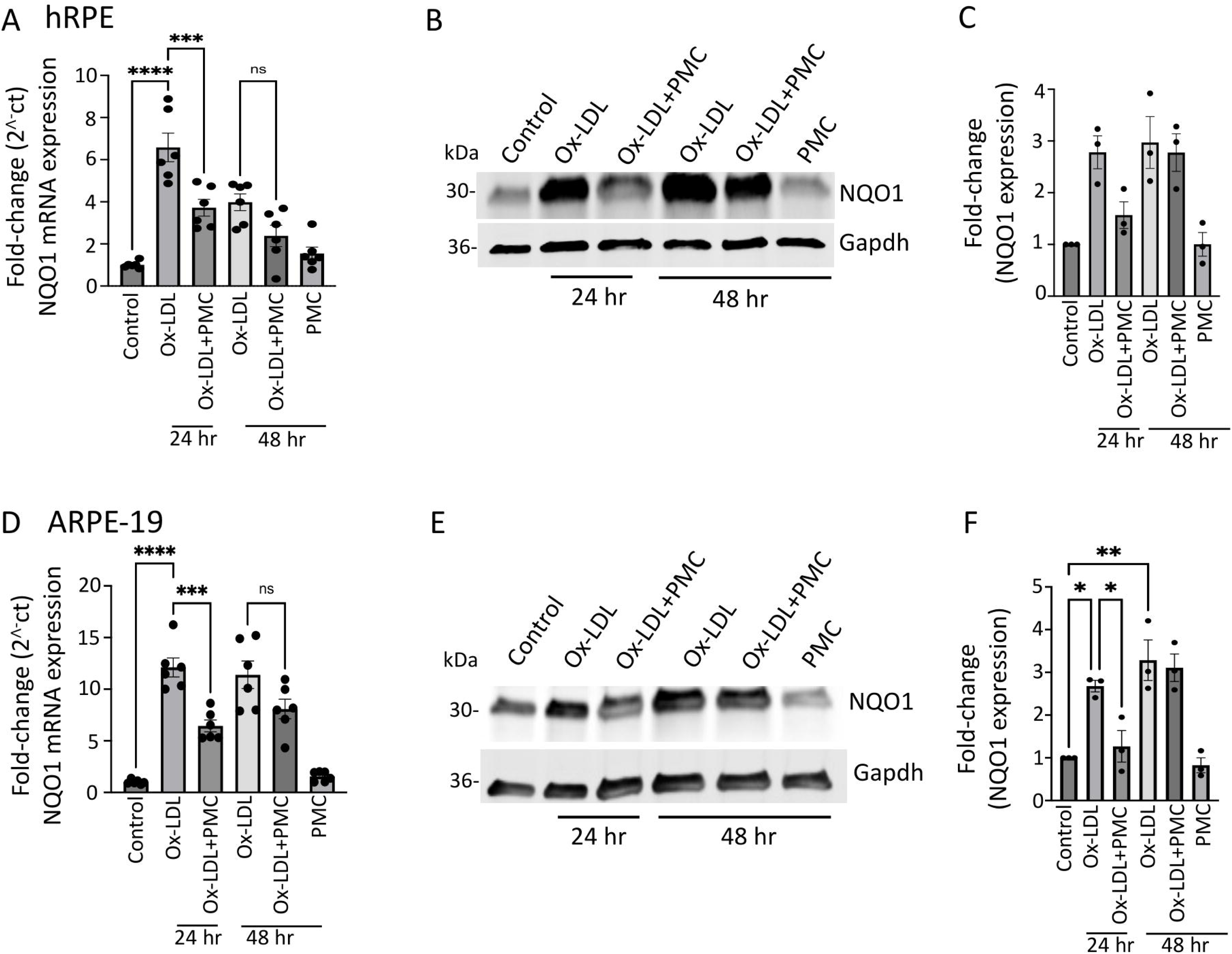
NQO1 upregulation in ox-LDL-treated RPE cells is suppressed by treatment with PMC. (A) Serum-starved hRPE cells were treated with ox-LDL (200 µg/ml) in the presence or absence of PMC (1.3 µM). NQO1 levels were determined using qPCR at 24 and 48 hr, n=6. (B) Western blot was conducted on cell lysates from hRPE cells treated with ox-LDL with or without PMC to determine NQO1 levels. (C) Densitometry was conducted with normalization for GAPDH, n=3. (D) Serum-starved ARPE-19 treated under the same experimental treatment conditions as above were analyzed for NQO1 levels using qPCR, n=6. (E) NQO1 protein levels were measured in cell lysates from ARPE-19 in the ox-LDL treated groups with or without PMC. (F) Densitometry analysis was conducted with normalization with GAPDH to quantify NQO1 levels, n=3. NQO1 levels were also measured in hRPE and ARPE-19 cells treated with PMC alone. Serum-starved untreated cells were considered as the control for all the experiments. Values are indicated as mean ± SEM of the indicated n. One-way ANOVA was used for statistical analysis, *P<0.05, **P<0.01, ***P<0.001

Similar results were obtained using ARPE-19 cells with a significant upregulation (12.1 ± 0.91-fold; p<0.001) of NQO1 mRNA at 24 hr in ox-LDL treated cells in comparison to control. Addition of PMC along with ox-LDL resulted in significantly lower levels of NQO1 mRNA induction at 24 hr (6.4 ± 0.6-fold; p<0.001, ox-LDL vs. ox-LDL+PMC). At 48 hr, there were 11.4 ± 1.3-fold and 8.1 ± 0.9-fold increases in NQO1 levels in cells treated with ox-LDL or ox-LDL plus PMC, respectively, compared to controls. PMC alone had no effect on NQO1 levels (Figure 3D). NQO1 proteins levels were significantly increased at 24 hr (2.7 ± 0.1-fold; p<0.05) and 48 hr (3.3 ± 0.5-fold; p<0.01) with ox-LDL treatments compared to controls. In contrast, a significantly lower level of NQO1 (1.3 ± 0.4-fold) was observed in ox-LDL and PMC treated cells at 24 hr compared to ox-LDL treated cells (p<0.05). At 48 hr, the level of NQO1 with ox-LDL+PMC treatment (3.1 ± 0.3-fold compared to control) was similar to that for ox-LDL treated cells (Figure 3E, F).

### PMC reduces ox-LDL-induced ROS

HO-1, an ER anchored protein, is known to play an important role during ER stress^28^. Thus, to investigate changes in intracellular ROS levels specifically in the ER, we treated ARPE-19 cells with ox-LDL, ox-LDL plus PMC or PMC alone for 24 hr and compared changes in ROS levels to control. Since the responses to ox-LDL and PMC by ARPE19 cells were confirmed to be comparable to those for hRPE cells in all experiments above, ARPE19 cells were used for subsequent mechanism of action experiments. Ox-LDL treatment resulted in elevated levels of ROS that were largely localized in the ER in comparison to control (Figure 4A). Quantification using Manders coefficient confirmed a significant colocalization of elevated ROS with ER in ox-LDL treated cells compared to control (0.9; p<0.001) (Figure 4B). Addition of PMC along with ox-LDL led to significantly lower levels of ROS generation as well as ROS/ER colocalization (0.8; p<0.01) compared to ox-LDL treated cells (Figure 4A and B). PMC alone had no effect on ROS levels or colocalization with ER compared to control (Figure 4A, B).

**Figure 4.**
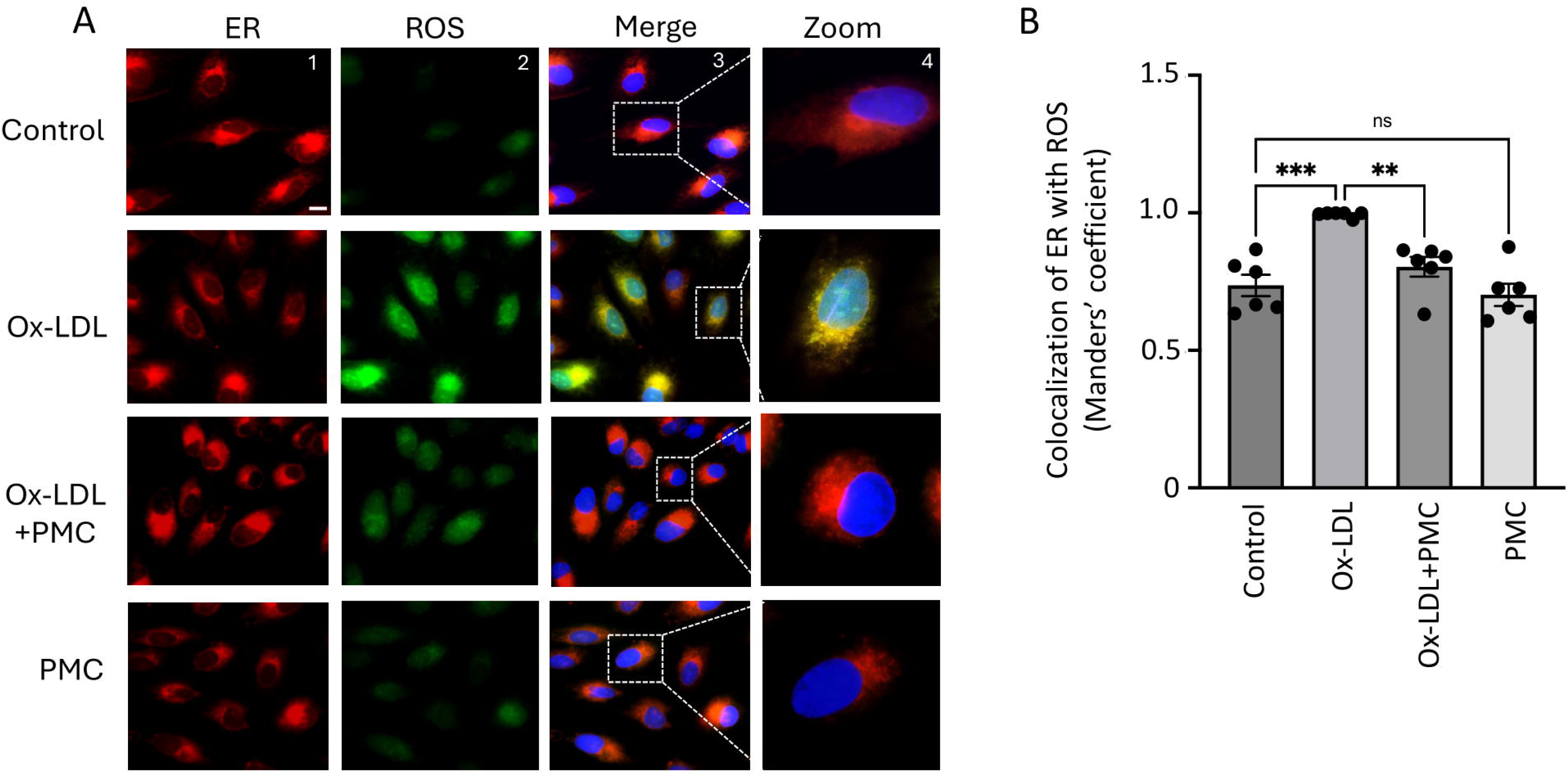
PMC prevents ox-LDL induced oxidative stress. (A) ARPE-19 cells were treated with ox-LDL (200 µg/ml) in the presence/absence of PMC (1.3 µM) for 24 hr. ROS levels were visualized using H2DCFDA stain (green) and localization of ROS in ER was estimated using the ER-Tracker™ (red). Changes in ROS levels in the ER were also examined in cells treated with PMC. Untreated cells were considered as controls. DAPI was used to stain the nucleus (blue), Scale bar= 10 µm. Panel 4 (zoom) are higher magnification images of the marked area in panel 3 (merge). (B) Quantification of colocalization of ROS with ER was measured using the Manders’ coefficient in ImageJ JaCoP plugin. Values are indicated as mean ± SEM of n=6. One-way ANOVA was used for statistical analysis, **P<0.01, ***P<0.001

### ox-LDL stimulates Nrf2 nuclear translocation

We next investigated ROS-mediated activation and nuclear translocation of Nrf2 because of its important role in antioxidant response in RPE. Results indicated upregulation of Nrf2 protein in the nucleus of cells treated with ox-LDL for 24 hr in comparison to control. This was further confirmed by a 3D reconstruction of the Z-stack confocal images of Nrf2 immunostaining (Figure 5A). Quantification of images using Manders coefficient confirmed a significant colocalization of Nrf2 with the nucleus in ox-LDL treated cells in comparison to control (0.57 v. 0.3; p<0.001) (Figure 5B). Cells treated with ox-LDL and PMC indicated lower levels of Nrf2 expression and a significant lower level of Nrf2 nuclear colocalization in comparison to ox-LDL treated cells (0.46 vs. 0.57; p<0.05) (Figure 5A and B). PMC treatment alone indicated similar levels of Nrf2 expression and nuclear colocalization as control (0.27 vs. 0.3).

**Figure 5.**
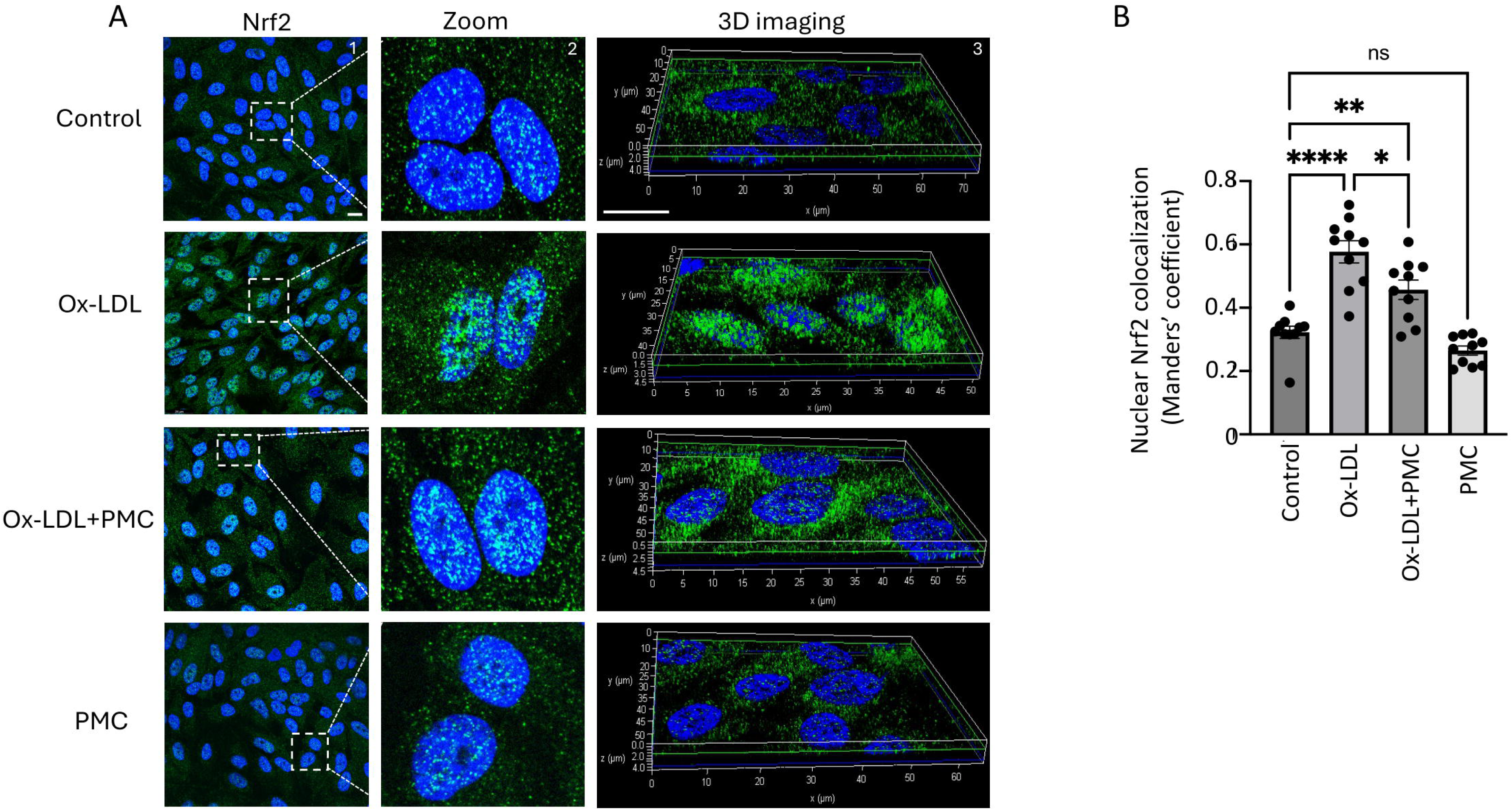
PMC prevents nuclear translocation of Nrf2. (A) Nuclear translocation of Nrf2 (green) was visualized in APRE-19 cells treated with ox-LDL (200 µg/ml) in the presence/absence of PMC (1.3 µM) for 24 hr. Nuclear translocation of Nrf2 was also imaged in PMC treated cells. Untreated cells were considered as the control. Nuclei were stained using DAPI (blue), Scale bar= 10 µm. Panel 2 (zoom) are higher magnification images of the marked area in panel 1 (Nrf2). Panel 3 shows the 3D reconstruction of the Z-stack images. (B) Quantification of the nuclear colocalization of Nrf2 was quantified using Manders’ coefficient in ImageJ JaCoP plugin. Values are indicated as mean ± SEM of n=10. One-way ANOVA was used for statistical analysis, *P<0.05, **P<0.01, ***P<0.001

Nrf2 nuclear translocation leads to activation of antioxidant response elements which activate HO-1. Since HO-1 is localized to the ER, we visualized the effects of ox-LDL treatment with and without PMC on HO-1 levels in the ER ^29^. Using the ER marker, calreticulin, we observed high HO-1 levels mainly in the ER of ARPE-19 cells treated with ox-LDL for 24 hr in comparison to control (Figure 6A). Quantification using the Manders coefficient confirmed the significantly elevated colocalization of HO-1 with calreticulin in the ER in comparison to control (0.9 vs. 0.6; p<0.001) (Figure 6B). Cells treated with ox-LDL and PMC had significantly lower levels of HO-1 and its colocalization with the calreticulin in the ER in comparison to ox-LDL alone (0.8 vs. 0.9; p<0.001) (Figure 6A, B). PMC had no effect on HO-1 expression levels or its colocalization with calreticulin, which was similar to that for the control (0.5 vs. 0.6) (Figure 6A, B).

**Figure 6.**
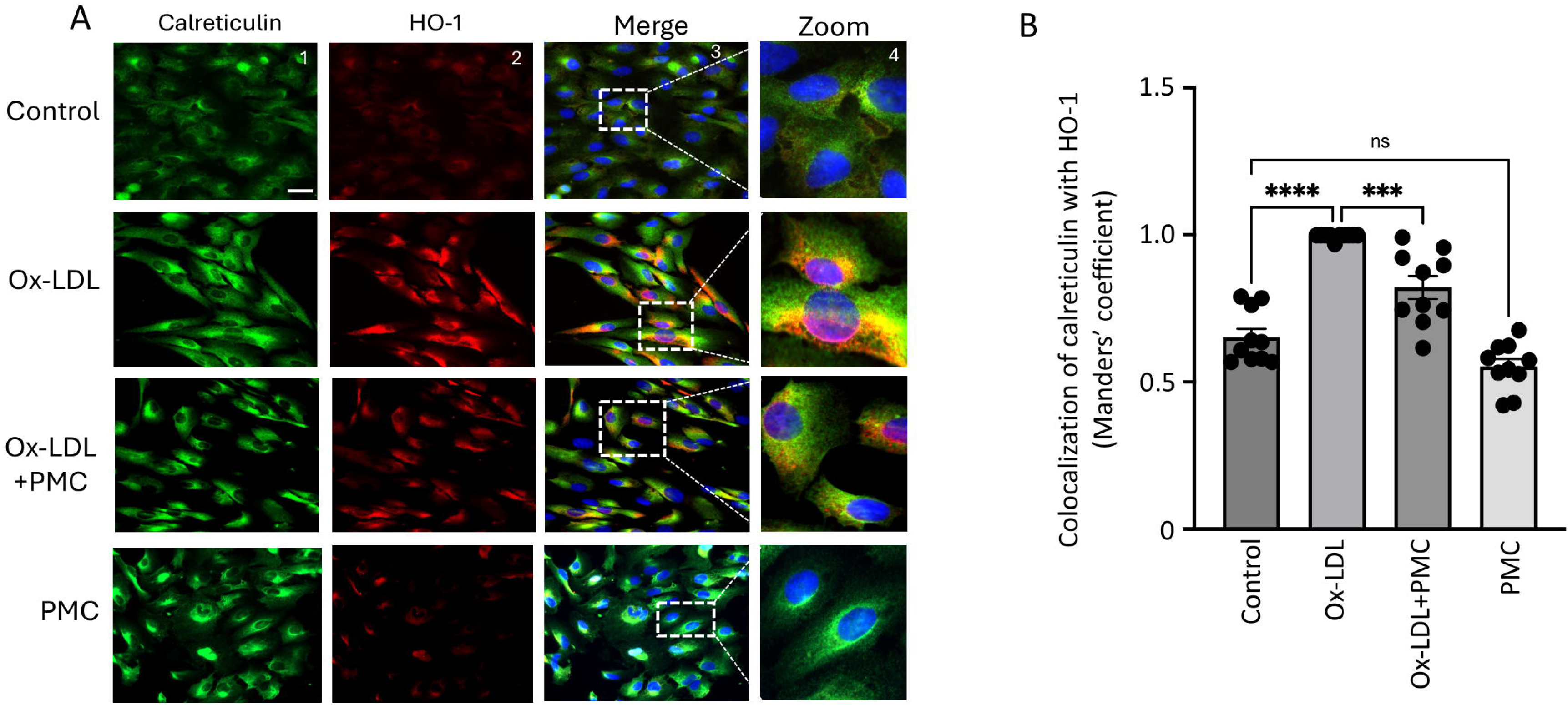
PMC prevents ox-LDL induced HO-1 upregulation. (A) ARPE-19 cells were treated with ox-LDL (200 µg/ml) with or without PMC (1.3 µM) for 24 hr. Following treatment, HO-1 (red) and ER localization calreticulin (green) were visualized. Nuclei were visualized using DAPI (blue), Scale bar= 25 µm. Panel 4 (zoom) is the higher magnification images of the marked area in panel 3. (B) Quantification of HO-1 colocalization with calreticulin was quantified based on Manders’ coefficient in ImageJ JaCoP plugin. Values are indicated as mean ± SEM of n=10. One-way ANOVA was used for statistical analysis, ***P<0.001

### PMC protects against ox-LDL via the HO-1 pathway

We next investigated the functional role of HO-1 for PMC-mediated cytoprotection against ox-LDL. This was accomplished by suppressing the expression of HMOX1 using siRNA (siHMOX1) for 48 hr followed by treatment with ox-LDL or ox-LDL plus PMC for an additional 24 hr in hRPE cells (Figure 7A). Conditioned media were collected for evaluation of cytotoxicity, and cell lysates were collected for protein analysis. HO-1 was undetectable in siRNA control (siScr) and siHMOX1 treated control cells (Figure 7B, C). Addition of ox-LDL for 24 hr led to significantly elevated HO-1 levels in siScr-treated control cells (28.6 ± 2.8-fold) which was significantly suppressed by siHMOX (2.0 ± 1.05 -fold; p<0.001 compared to siScr), confirming highly efficient knockdown of siHMOX1 expression. Treatment with ox-LDL and PMC for 24 hr led to significantly lower levels of HO-1 in siScr cells in comparison to siScr cells treated with ox-LDL (13.5 ± 2.3-fold vs. 28.6 ± 2.8-fold; p<0.001). siHMOX1 silenced cells treated with ox-LDL and PMC had significantly lower levels of HO-1 in comparison to siScr cells treated with ox-LDL plus PMC (1.1 ± 0.1-fold vs. 13.5 ± 2.3-fold; p<0.01), further confirming HMOX1 expression silencing at the protein level (Figure 7B, C).

**Figure 7.**
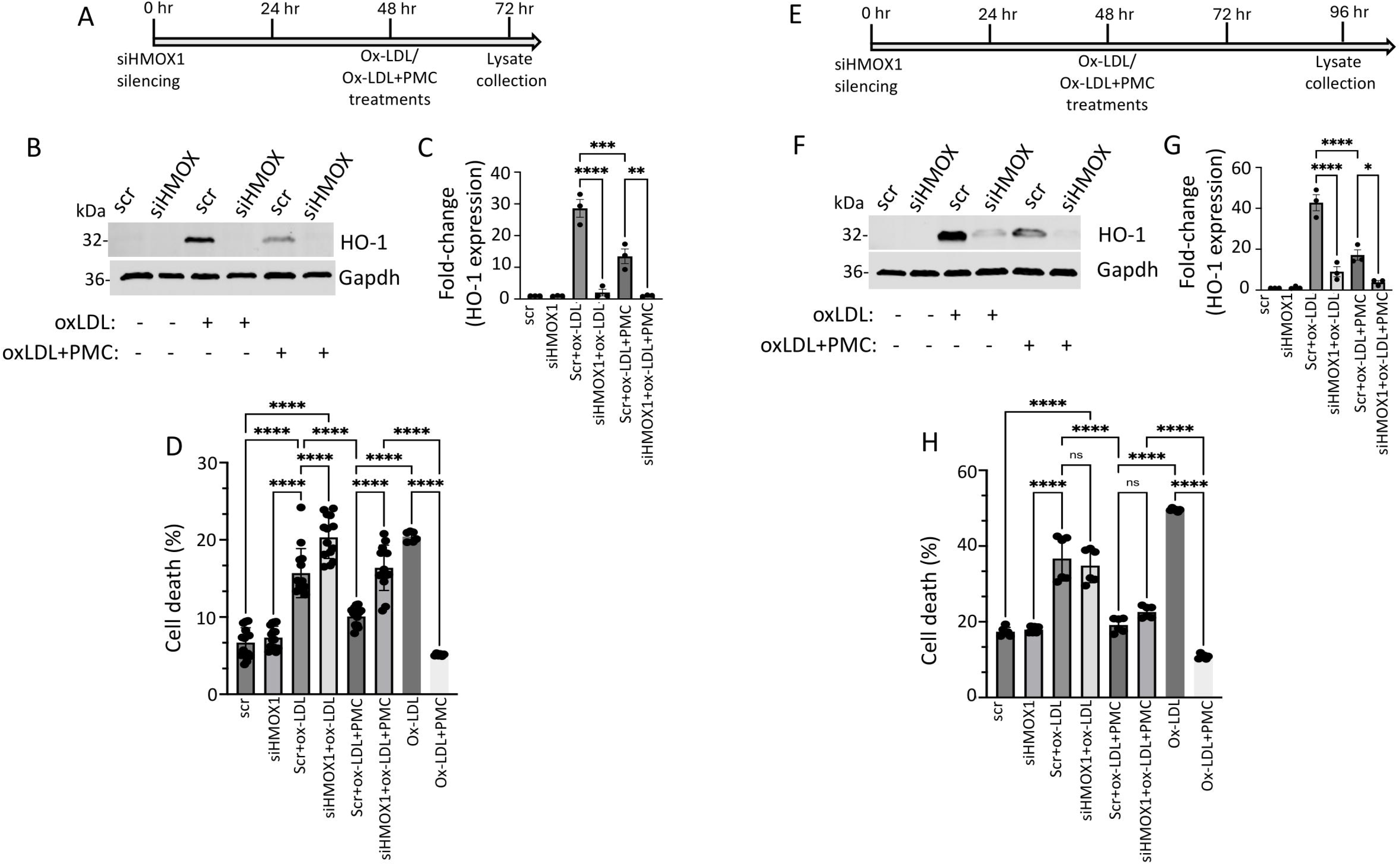
HO-1 contributes to PMC protection against ox-LDL. (A) Illustration showing the timeline of 48 hr siHMOX1 silencing and 24 hr ox-LDL treatments in the presence or absence of PMC in hRPE cells. (B) Western blotting was conducted to estimate HO-1 levels in hRPE cells subjected to siScr or siHMOX1 for 48 hr followed by ox-LDL (200 µg/ml) treatments with or without PMC (1.3 µM). (C) Quantification of HO-1 levels was performed using densitometry after normalization with GAPDH, n=3. (D) LDH levels were measured in the conditioned media from siScr or siHMOX-treated cells along with the siScr and siHMOX1 cells that were treated with ox-LDL with or without PMC, n=14. (E) Illustration of the timeline indicating, 48 hr siHMOX1 silencing in hRPE cells followed by additional 48 hr treatment of ox-LDL with/without PMC. (F) HO-1 levels were detected in hRPE cells treated with siScr or siHMOX1 for 48 hr, followed by ox-LDL in the presence or absence of PMC using western blotting. (G) Densitometry analysis was conducted to quantify HO-1 levels after normalization with GAPDH, n=3. (H) Cytotoxicity was analyzed by assaying LDH in the conditioned media of siScr and siHMOX1 and siScr and siHMOX cells that were subjected to ox-LDL with/without PMC treatment, n=6. Cells treated with ox-LDL in the presence or absence of PMC without siScr or siHMOX1 were used as control for both 24 hr and 48 hr ox-LDL and ox-LDL with PMC treatments, n=6. Values are indicated as mean ± SEM of the indicated n. One-way ANOVA was used for statistical analysis, *P<0.05, **P<0.01, ***P<0.001

Cytotoxicity analysis of cells treated with ox-LDL without siScr or siHMOX1 displayed 20.4 ± 0.3% cell death, which was significantly reduced by PMC to 5 ± 0.1% (p<0.001). Treatment of hRPE cells with siHMOX1 resulted in a very mild cytotoxicity effect which is comparable to that for siScr control (7.3 ± 0.4% vs. 6.7 ± 0.5%). Introduction of ox-LDL to siScr-treated (control) cells led to a significant increase in the cell death compared to siScr control (15.7 ± 0.8 % vs. 6.7 ± 0.5%; p<0.001). The addition of ox-LDL to siHMOX1 cells resulted in a further significant increase in the cell death compared to siScr cells treated with ox-LDL (20.3 ± 0.7%. vs. 15.7 ± 0.8 %, p<0.001), indicating that HMOX1 is involved in the protection of cells from ox-LDL induced cytotoxicity. To assess the therapeutic efficacy of PMC in the presence and absence of HMOX1, cell death was measured in siScr cells and siHMOX1 cells treated with ox-LDL plus PMC. Results indicated 10.1 ± 0.3% cell death in siScr cells treated with ox-LDL plus PMC, which was significantly (p<0.001) lower than in siScr cells treated ox-LDL cells. Silencing of HMOX1 in cells treated with ox-LDL and PMC showed a reduced protection of cells at 24 hr compared to siScr cells with ox-LDL plus PMC (16.4 ± 0.8% v. 10.1 ± 0.3% cell death; p<0.001) (Figure 7D), confirming the role of HO-1 in PMC-mediated protection of RPE cells against ox-LDL. Comparison of % cell death induced by ox-LDL in all treatment conditions showed that suppressions of HMOX1 were associated with worsening of cell death with and without PMC treatment (Figure 7D).

Similar studies for the effects of HMOX1 silencing in PMC-mediated protection against ox-LDL were conducted with hRPE cells that were treated for 48 hr with ox-LDL with and without PMC (Figure 7E). Similar to hRPE cells at 24 hr (Figure 7B), siHMOX1 treated cells at 48 hr displayed very low levels of HO-1 (96 hr post siRNA). Addition of ox-LDL to siScr cells led to a robust increase in HO-1 while siHMOX1 significantly reduced HO-1 levels in cells with ox-LDL treatment (42.8 ± 3.9-fold vs. 9 ± 2.5-fold; p<0.001) (Figure 7F, G). Treatment of siScr cells with ox-LDL plus PMC led to a 17.2 ± 2.6-fold increase in HO-1 levels, which was significantly (p<0.001) lower than that in siScr cells treated with ox-LDL alone at 42.8 ± 3.9-fold. Additionally, siHMOX1 cells treated with ox-LDL plus PMC displayed significantly lower levels of HO-1 compared to siScr cells treated with ox-LDL plus PMC (3.8 ± 0.8-fold vs. 17.2 ± 2.6-fold; p<0.05) (Figure 7F, G). Furthermore, siHMOX1 treated cells at 48 hr indicated slightly higher levels of HO-1 with ox-LDL (9 ± 2.5-fold) and ox-LDL plus PMC (3.8 ± 0.8-fold) treated cells in comparison to 24 hr treatment of ox-LDL (2.1 ± 1.1-fold) and ox-LDL plus PMC (1.1 ± 0.1-fold) treated cells (Figure 7B, C vs. 7F, G), confirming the siRNA-mediated knockdown of HO-1 in primary hRPE cells lasted up to 96 hrs.

Cell death in response to 48 hr of ox-LDL treatment with and without HO-1 knockdown, and with and without PMC was measured in cells using the LDH cytotoxicity assay. Cells treated with ox-LDL with and without PMC, and with and without HMOX1 silencing were used as controls. As expected, cells treated for 48 hr with PMC and ox-LDL exhibited significantly less cell death compared to ox-LDL alone (10.8 ± 0.3% vs. 49 ± 0.1%; p<0.001). There were similar levels of cell death in siScr (17 ± 0.5%) and siHMOX1 (18 ± 0.3%) cells. Ox-LDL treatment of siScr cells led to comparable levels of cell death as for siHMOX1 cells with ox-LDL treatment (36.7 ± 2.4% vs. 34.8 ± 1.8%) (Figure 7H). The inclusion of PMC to ox-LDL treated siScr cells led to significantly less cell death in comparison to siScr cells treated with ox-LDL (19.0 ± 0.8% vs. 36.7 ± 2.4%; p<0.001). Interestingly, in contrast to 24 hr treatment, 48 hr treatment with ox-LDL plus PMC in siHMOX1 cells resulted in comparable levels of cell death to siScr cells treated with ox-LDL plus PMC (22.6 ± 0.6% vs. 19.0 ± 0.8%; p>0.05) (Figure 7H). This observation at 48 hr further corroborated the role of HO-1 in PMC mediated protection of cells against ox-LDL.

### Continuous presence of PMC provides optimal protection of RPE against ox-LDL

Concurrent treatment of PMC with ox-LDL prevented ox-LDL induced cellular death by over 80%; ox-LDL treatment for 48 hr induced significant cell death at 46.7 ± 0.1% compared to untreated control cells at 7.4 ± 0.1% (p<0.001), while simultaneous ox-LDL and PMC treatment significantly decreased cell death to the control levels at 7.0 ± 0.1% compared to ox-LDL alone (p<0.001) (Figure 8A). Next, we investigated the effects of PMC-mediated protection under different treatment conditions and durations. Pretreatment of hRPE with PMC for 24 hr followed by complete replacement of the media with ox-LDL plus PMC treatment for an additional 48 hr resulted in a high level of cellular protection (8.2 ± 0.1% cell death) as seen with 48 hr of simultaneous ox-LDL and PMC treatment (Figure 8A). Pretreatment of hRPE cells with PMC for 24 hr followed by complete replacement with media containing ox-LDL for an additional 48 hr did not provide any cellular protection (45.3 ± 1.2% cell death), which was similar to that observed with 48 hr of ox-LDL treatment alone. Exposure of cells pre-treated with PMC (24 hr) and addition of ox-LDL in the same media without media change led to a significant protection of cells at 48 hr as compared to ox-LDL treated cells (15.7 ± 0.2% vs. 46.7 ± 0.1%; p<0.001). Together, the data indicated that the continuous presence of PMC provides optimal protection of cells against ox-LDL, suggesting that PMC likely has short pharmacokinetics in RPE cells.

**Figure 8.**
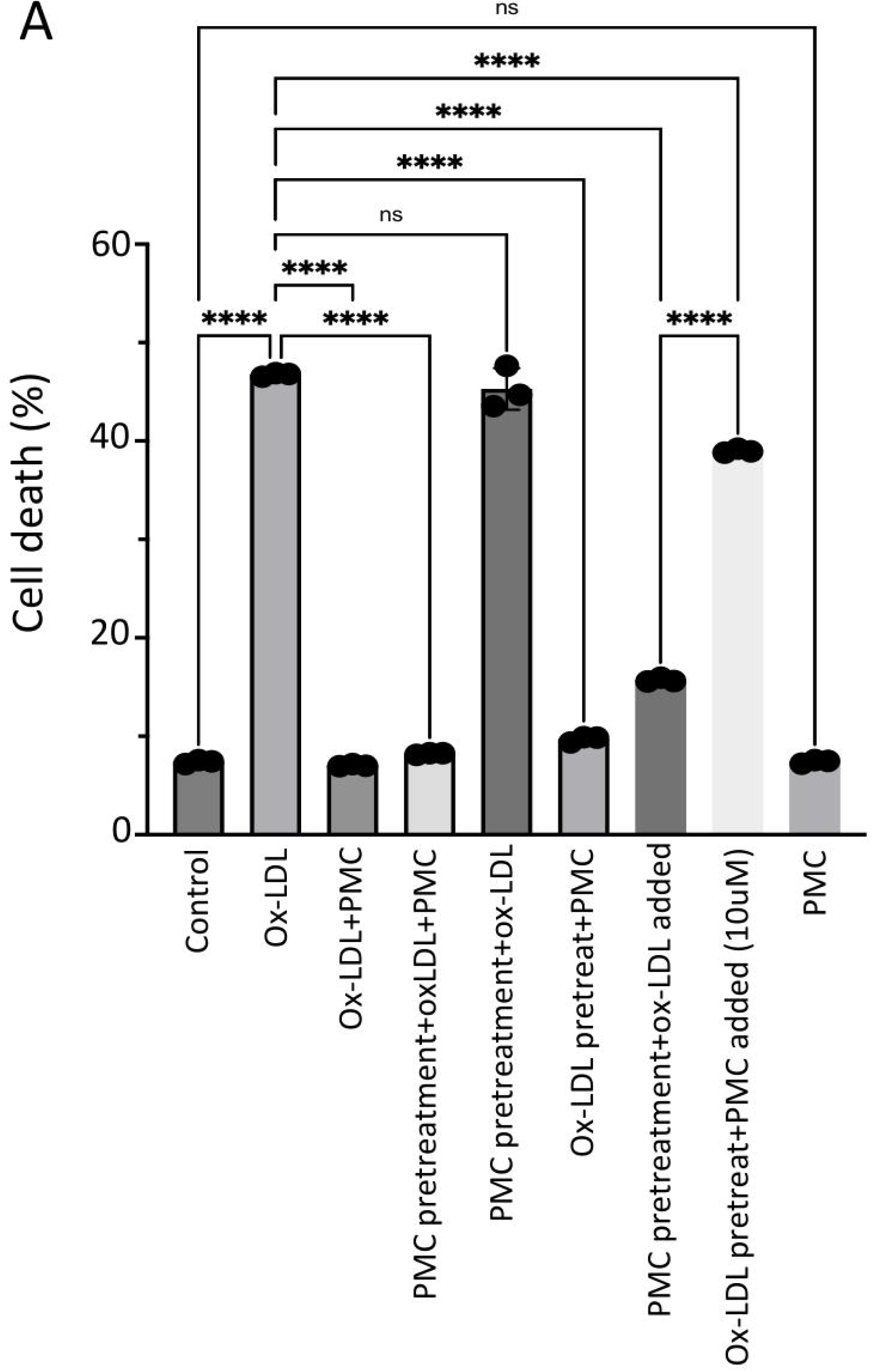
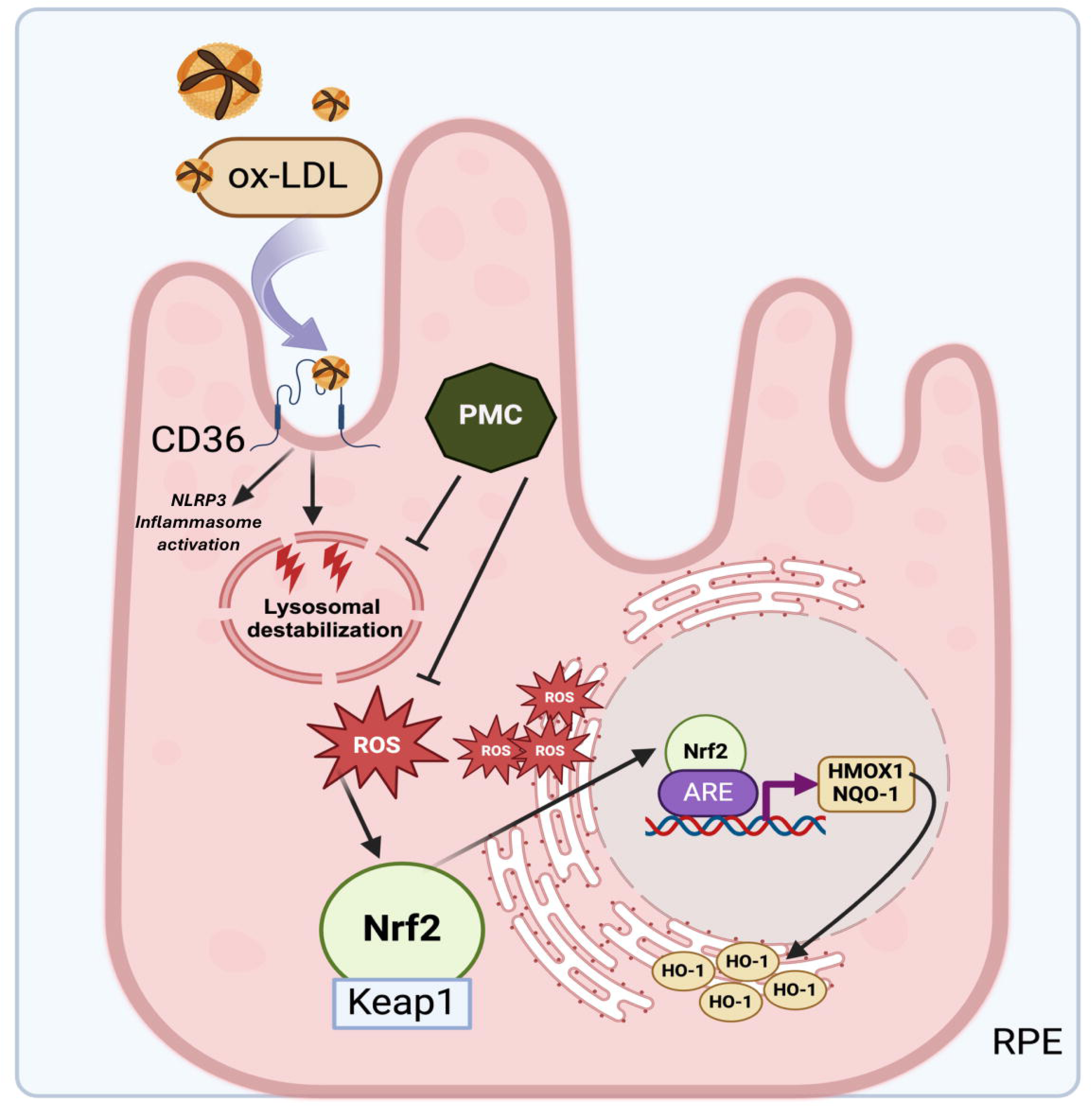
Continuous PMC presence is required for protection against ox-LDL. (A) Serum-starved hRPE cells were treated with either ox-LDL (200 µg/ml) and PMC (1.3 µM) alone or, ox-LDL and PMC under different treatment conditions. ox-LDL+PMC denotes simultaneous treatment, while in other treatment groups either the cells were pretreated with PMC or with ox-LDL. Following these treatments, media were replaced with ox-LDL+PMC, ox-LDL alone, or PMC alone. In another group, media were not replaced, and treatments were added to the same media. Cells were pretreated with PMC and ox-LDL was added to the same media or pretreated with ox-LDL and an approximately 10-fold higher PMC concentration (10 µM) was added to the same media. Untreated cells were considered as controls. Values are indicated as mean ± SEM of n=3. One-way ANOVA was used for statistical analysis, ***P<0.001. (B) Graphical summary of the PMC mediated protection against ox-LDL in RPE. Uptake of ox-LDL via the CD36 receptor^25^ causes lysosomal destabilization^26^ and oxidative stress in RPE leading to ROS generation. This triggers Nrf2 dissociation from Keap1 and its translocation to the nucleus where it interacts with the specific promoter region, ARE, leading to the upregulation of HMOX1 and NQO1. PMC prevents this upregulation of HMOX1/HO-1 and NQO1 levels by preventing ROS generation. Created using BioRender.

Pre-treatment of cells with ox-LDL for 48 hr to allow some cellular uptake of ox-LDL, followed by complete replacement with media containing PMC provided a significant cellular protection compared to 48 hr of ox-LDL treated cells (9.7 ± 0.2% vs. 46.7 ± 0.1% cell death; p<0.001) (Figure 8A). However, pre-treatment with ox-LDL for 48 hr followed by the addition of a high concentration of PMC at 10µM to the same culture media although provided a significant protection compared to ox-LDL treated cells (39 ± 0.1% vs. 46.7 ± 0.1%; p<0.001), there was significantly more cell death compared to PMC pretreated cells with ox-LDL added to the same media (39 ± 0.1% vs. 15.7 ± 0.1%; p<0.05). These results indicated that PMC is not very effective in preventing cell death for cells that are severely damaged from pro-longed (>48 hr) and continuous exposure to ox-LDL without PMC in this model. PMC treatment alone had no effect on cell viability (7.4 ± 0.1% cell death), which was similar to control (7.4 ± 0.1%) (Figure 8A).

## Discussion

AMD is a retinal degenerative disease, characterized by the formation and accumulation of drusen^30^. Lipids, including lipofuscin and phospholipids of phagocytized rod outer segments, are the major components of drusen^31^ and are prone to oxidation. Drusen expansion and coalescence lead to RPE dysfunction/atrophy and photoreceptor cell death, which are the hallmarks of AMD-associated pathologies^32,33^. Numerous studies have confirmed the presence of significantly higher levels of oxidized phospholipids in the macular^6^ and plasma^34^ of AMD patients, supporting the correlation between oxidized lipids exposure and AMD pathogenesis.

Previous studies in our lab have shown the uptake of ox-LDL through the CD36 receptor and a dose-dependent cytotoxicity of ox-LDL in RPE cells^25^. In line with this, our lab reported the significant role of NLRP3 inflammasome activation as the major contributor of cytokine release and cell death in ox-LDL treated RPE cells^25^. Based on these observations, ox-LDL induced damage to RPE cells was used as a model in the present study to mimic AMD associated pathologies in RPE cells to determine the mechanism of PMC-mediated RPE protection against ox-LDL. Using this model, our study unraveled the key antioxidant pathway through which PMC protects RPE cells from oxidative damage and cytotoxicity induced by ox-LDL. We show the efficacy of PMC in preventing ox-LDL–induced ROS generation thus, preventing activation of Nrf2-ARE-HMOX1/NQO1 pathway which gets activated during ox-LDL induced oxidative damage.

We have previously screened multiple antioxidants against ox-LDL induced cytotoxicity. Our results revealed the significant efficacy of sterically hindered phenol compounds in protecting cells from oxidative stress by scavenging free radicals and preventing ox-LDL induced lysosomal destabilization and death in RPE cells^26^. Consistent with this, a sterically hindered phenol, PMC presented high antioxidant and protective efficacy in RPE cells against ox-LDL and was thus, used in our present study.

One of the primary pathway employed by RPE to maintain cellular homeostasis and neutralize oxidative stress is the Nrf2 pathway^14^. Nrf2, a master transcriptional factor under oxidative stimuli and ROS generation, dissociates from Keap1 and translocates to the nucleus. The subsequent signaling cascade causes Nrf2 nuclear translocation and interaction with the specific ARE promoter region leads to the upregulation of antioxidant genes including HMOX1 and NQO1^18,35^. Additionally, HO-1 proteins that are located on the endoplasmic reticulum (ER) are also released from the ER to the cytosol under oxidative stress^18^.

Due to the antioxidant and anti-inflammatory properties of HO-1, it was considered as a promising therapeutic strategy against age-related diseases of the eye. However, emerging evidence suggested also deleterious functions of HO-1 since overexpression of HO-1 has been linked with excessive Fe^2+^ and ROS generation, which can lead to lipid peroxidation and ferroptosis, an iron-dependent cell death pathway which is a critical hallmark of AMD^20^. Previously, a study showed that HO-1 upregulation in a NaIO3 induced AMD mouse model can lead to elevated iron levels which in turn causes free radical-mediated damage^23^. Interestingly, knockdown of HO-1 using siRNA reduced oxidative stress induced ferroptosis in NaIO3 treated ARPE-19 cells. Moreover, suppression of HO-1 levels using a known inhibitor, Zn-protoporphyrin (ZnPP), recused RPE degeneration, restored visual function and retinal structure in the NaIO3 AMD mice model^23^. In another study, knockdown of HO-1 demonstrated reduced proliferation and migration of endothelial cells suggesting it to be a potential target in treating choroidal neovascularization that occurs in advanced neovascular or wet AMD^36^.

Previous oxidative stress models of AMD have reported a similar upregulation of HO-1 as observed in our study. Cigarette smoke extract, an important environmental risk factor for AMD indicated upregulation of HO-1 and a modest involvement of Nrf2^16^. A H2O2 induced ROS production model of AMD confirmed the upregulation of HO-1 and NQO1 in human ARPE-19 cells^17^. These observations are in line with our observed HO-1 upregulation with ox-LDL treated cells.

Interestingly, concurrent treatment of ox-LDL with PMC, resulted in lower levels of HO-1 and NQO1 in both hRPE and ARPE-19 cells. One of the major contributors for this could be the role of PMC in preventing the upstream oxidative damage caused by internalized ox-LDL in cells^25^, which prevents Nrf2 nuclear translocation and subsequent upregulation of HO-1 and NQO1. Notably, the transcriptional and translational changes for HO-1 and NQO1 were comparable in both hRPE and ARPE-19, validating ARPE-19 as a useful model to understand the cellular mechanism involved in PMC-mediated protection from ox-LDL induced pathologies. The relevance of ARPE-19 cells as a model in the discovery of innovative therapeutic interventions has been witnessed earlier^37,38^.

Over-activation of HO-1 has previously been linked to retinal degeneration through ER stress. It was previously observed that low levels of HO-1 provided protection against light-induced photoreceptor cell death while high levels of HO-1 induced photoreceptor cell death and retinal degeneration^39^. Consistent with this, we observed significantly higher levels of HO-1 with ox-LDL treatments in comparison to ox-LDL and PMC concurrent treatment. Based on this, we validated the efficacy of PMC in preventing ROS, and subsequent activation of Nrf2-ARE-HMOX1 axis. Notably, our data indicated that suppression of HMOX1 expression by siRNA resulted in exacerbated ox-LDL induced cell death. Importantly, this effect was also observed in cells concurrently treated with ox-LDL and PMC, indicating cellular protection depends on the maintenance of optimal levels of HO-1 levels. The complete absence of HO-1 hinders the intrinsic cellular defense against oxidative stress. Our data also showed that PMC provides protection from ox-LDL at least partly involves the Nrf2-ARE-HMOX1 pathway, since the protection by PMC against ox-LDL in RPE cells was significantly reduced upon suppression of HMOX1 expression by siRNA.

While there are a number of ongoing research into antioxidant therapies for AMD^33,40-42^, sterically hindered phenols could be an attractive candidate due to their stability and ability to protect retinal cells from oxidative stress induced by ox-LDL, a key driver in AMD pathology. Future investigations in our lab will focus on identifying and validating multiple different pathways involved in PMC-mediated protection and defining the role of additional signaling cascades that may contribute to its therapeutic potential.

## Data statement

Datasets and materials used in this study are available upon request from the corresponding author

## Acknowledgments

We thank Xinyao Hu at the Schepens Eye Research Institute Morphology Core for assistance with the confocal microscope.

## Funding

This work was supported by Massachusetts Lions Eye Research fund, Grimshaw-Gudewicz Foundation AMD research grant, Gilbert Family Foundation, and NIH National Eye Institute core grant P30EYE003790.

## Author contributions

Writing: SC, YEN, PAD, Conceptualization: SC, YEN, PAD, Data curation: SC, JM, Methodology, investigation: SC, JM, ZH, Analysis: SC, JM, EK, SP, PYB, Writing editing: SC, JM, ZH, EK, YEN, PAD.

## Notes

### Competing Interest Statement

The authors have declared no competing interest.

## References

1 Sacconi, R., Corbelli, E., Querques, L., Bandello, F. & Querques, G. A review of current and future management of geographic atrophy. Ophthalmology and therapy 6, 69–77 (2017).

2 Ebrahimi, K. B. & Handa, J. T. Lipids, lipoproteins, and age-related macular degeneration. Journal of lipids 2011, 802059 (2011).

3 Basyal, D., Lee, S. & Kim, H. J. Antioxidants and Mechanistic Insights for Managing Dry Age-Related Macular Degeneration. Antioxidants 13, 568 (2024).

4 Evereklioglu, C. et al. Nitric oxide and lipid peroxidation are increased and associated with decreased antioxidant enzyme activities in patients with age-related macular degeneration. Documenta ophthalmologica 106, 129–136 (2003).

5 Abokyi, S., To, C.-H., Lam, T. T. & Tse, D. Y. Central role of oxidative stress in age-related macular degeneration: evidence from a review of the molecular mechanisms and animal models. Oxidative medicine and cellular longevity 2020, 7901270 (2020).

6 Suzuki, M. et al. Oxidized phospholipids in the macula increase with age and in eyes with age-related macular degeneration. Molecular vision 13, 772 (2007).

7 Kamei, M. et al. Scavenger receptors for oxidized lipoprotein in age-related macular degeneration. Investigative ophthalmology & visual science 48, 1801–1807 (2007).

8. Xue, C. C., et al. Lipid-lowering drug and complement factor H genotyping–personalized treatment strategy for age-related macular degeneration. iScience 27 (2024).

9 Batliwala, S., Xavier, C., Liu, Y., Wu, H. & Pang, I.-H. Involvement of Nrf2 in ocular diseases. Oxidative medicine and cellular longevity 2017, 1703810 (2017).

10 Study, T. A.-R. E. D. The age-related eye disease study (AREDS): Design implications AREDS report no. 1. Controlled clinical trials 20, 573-600 (1999).

11 Group, A.-R. E. D. S. R. Lutein+ zeaxanthin and omega-3 fatty acids for age-related macular degeneration: the Age-Related Eye Disease Study 2 (AREDS2) randomized clinical trial. Jama 309, 2005–2015 (2013).

12 Keenan, T. D. et al. Oral antioxidant and lutein/zeaxanthin supplements slow geographic atrophy progression to the fovea in age-related macular degeneration. Ophthalmology (2024).

13 Sachdeva, M. M., Cano, M. & Handa, J. T. Nrf2 signaling is impaired in the aging RPE given an oxidative insult. Experimental eye research 119, 111–114 (2014).

14 Lambros, M. L. & Plafker, S. M. Oxidative stress and the Nrf2 anti-oxidant transcription factor in age-related macular degeneration. Retinal Degenerative Diseases: Mechanisms and Experimental Therapy, 67–72 (2016).

15 Catanzaro, M. et al. Eye-light on age-related macular degeneration: Targeting Nrf2-pathway as a novel therapeutic strategy for retinal pigment epithelium. Frontiers in Pharmacology 11, 844 (2020).

16 Bertram, K. M., Baglole, C. J., Phipps, R. P. & Libby, R. T. Molecular regulation of cigarette smoke induced-oxidative stress in human retinal pigment epithelial cells: implications for age-related macular degeneration. American Journal of Physiology-Cell Physiology 297, C1200–C1210 (2009).

17 Muangnoi, C., Sharif, U., Ratnatilaka Na Bhuket, P., Rojsitthisak, P. & Paraoan, L. Protective effects of curcumin ester prodrug, curcumin diethyl disuccinate against H2O2-induced oxidative stress in human retinal pigment epithelial cells: potential therapeutic avenues for age-related macular degeneration. International Journal of Molecular Sciences 20, 3367 (2019).

18 Liu, R. et al. Heme oxygenase 1 in erythropoiesis: an important regulator beyond catalyzing heme catabolism. Annals of Hematology 102, 1323–1332 (2023).

19 Kikuchi, G., Yoshida, T. & Noguchi, M. Heme oxygenase and heme degradation. Biochemical and biophysical research communications 338, 558–567 (2005).

20 Wei, D. et al. The significance of precisely regulating heme oxygenase-1 expression: Another avenue for treating age-related ocular disease? Ageing Research Reviews, 102308 (2024).

21 Shin, C. Y. & Jeong, K. W. Photooxidation of A2E by blue light regulates heme oxygenase 1 expression via NF-κB and lysine methyltransferase 2A in ARPE-19 cells. Life 12, 1698 (2022).

22 Yang, P.-M., Cheng, K.-C., Yuan, S.-H. & Wung, B.-S. Carbon monoxide-releasing molecules protect against blue light exposure and inflammation in retinal pigment epithelial cells. International Journal of Molecular Medicine 46, 1096–1106 (2020).

23 Tang, Z. et al. HO-1-mediated ferroptosis as a target for protection against retinal pigment epithelium degeneration. Redox biology 43, 101971 (2021).

24 Chen, C. et al. Induction of ferroptosis by HO-1 contributes to retinal degeneration in mice with defective clearance of all-trans-retinal. Free Radical Biology and Medicine 194, 245–254 (2023).

25 Gnanaguru, G., Choi, A. R., Amarnani, D. & D’Amore, P. A. Oxidized lipoprotein uptake through the CD36 receptor activates the NLRP3 inflammasome in human retinal pigment epithelial cells. Investigative ophthalmology & visual science 57, 4704–4712 (2016).

26 Gnanaguru, G. et al. Discovery of sterically-hindered phenol compounds with potent cytoprotective activities against ox-LDL–induced retinal pigment epithelial cell death as a potential pharmacotherapy. Free Radical Biology and Medicine 178, 360–368 (2022).

27 Fernandes, M., McArdle, B., Schiff, L. & Blenkinsop, T. A. Stem cell–derived retinal pigment epithelial layer model from adult human globes donated for corneal transplants. Current Protocols in Stem Cell Biology 45, e53 (2018).

28 Gáll, T., Balla, G. & Balla, J. Heme, heme oxygenase, and endoplasmic reticulum stress—a new insight into the pathophysiology of vascular diseases. International journal of molecular sciences 20, 3675 (2019).

29 Kim, H. P. et al. Heme oxygenase-1 comes back to endoplasmic reticulum. Biochemical and biophysical research communications 404, 1–5 (2011).

30 Kaarniranta, K. et al. Autophagy in age-related macular degeneration. Autophagy 19, 388–400 (2023).

31 Handa, J. T., Cano, M., Wang, L., Datta, S. & Liu, T. Lipids, oxidized lipids, oxidation-specific epitopes, and Age-related Macular Degeneration. Biochimica et Biophysica Acta (BBA)-Molecular and Cell Biology of Lipids 1862, 430–440 (2017).

32 Sparrow, J. R. & Boulton, M. RPE lipofuscin and its role in retinal pathobiology. Experimental eye research 80, 595–606 (2005).

33 Kushwah, N., Bora, K., Maurya, M., Pavlovich, M. C. & Chen, J. Oxidative stress and antioxidants in age-related macular degeneration. Antioxidants 12, 1379 (2023).

34 Ikeda, T. et al. Paraoxonase gene polymorphisms and plasma oxidized low-density lipoprotein level as possible risk factors for exudative age-related macular degeneration. American journal of ophthalmology 132, 191–195 (2001).

35 Wu, J. et al. The non-canonical effects of heme oxygenase-1, a classical fighter against oxidative stress. Redox biology 47, 102170 (2021).

36 Zhang, W. et al. Silencing heme oxygenase-1 gene expression in retinal pigment epithelial cells inhibits proliferation, migration and tube formation of cocultured endothelial cells. Biochemical and Biophysical Research Communications 434, 492–497 (2013).

37 Peterson, K. M. et al. Serum-deprivation response of ARPE-19 cells; expression patterns relevant to age-related macular degeneration. Plos one 19, e0293383 (2024).

38 Mishra, S. et al. Accumulation of cholesterol and increased demand for zinc in serum-deprived RPE cells. Molecular vision 22, 1387 (2016).

39 Li, H. et al. High dose expression of heme oxigenase-1 induces retinal degeneration through ER stress-related DDIT3. Molecular neurodegeneration 16, 1–17 (2021).

40 Kulbay, M., Wu, K. Y., Nirwal, G. K., Bélanger, P. & Tran, S. D. The Role of Reactive Oxygen Species in Age-Related Macular Degeneration: A Comprehensive Review of Antioxidant Therapies. Biomedicines 12, 1579 (2024).

41 Wong, K.-H. et al. Discovering the potential of natural antioxidants in age-related macular degeneration: a review. Pharmaceuticals 15, 101 (2022).

42 Evans, J. Antioxidant vitamin and mineral supplements for age-related macular degeneration. The Cochrane database of systematic reviews, CD000254- (2002).

